# Hsf1 and Hsp70 constitute a two-component feedback loop that regulates the yeast heat shock response

**DOI:** 10.1101/183137

**Authors:** Joanna Krakowiak, Xu Zheng, Nikit Patel, Jayamani Anandhakumar, Kendra Valerius, David S. Gross, Ahmad S. Khalil, David Pincus

**Affiliations:** Whitehead Institute for Biomedical Research, Cambridge, USA; Department of Biomedical Engineering and Biological Design Center, Boston University, Boston, USA; Wyss Institute for Biologically Inspired Engineering, Harvard University, Boston, USA; Department of Biochemistry and Molecular Biology, Louisiana State University Health Sciences Center, Shreveport, USA

## Abstract

Models for regulation of the eukaryotic heat shock response typically invoke a negative feedback loop consisting of the transcriptional activator Hsf1 and a molecular chaperone encoded by an Hsf1 target gene. Previously, we identified Hsp70 as the chaperone responsible for Hsf1 repression in *Saccharomyces cerevisiae* and constructed a mathematical model based on Hsp70-mediated negative feedback that recapitulated the dynamic activity of Hsf1 during heat shock. The model was based on two assumptions: dissociation of Hsp70 activates Hsf1, and transcriptional induction of Hsp70 deactivates Hsf1. Here we validated these assumptions. First, we severed the feedback loop by uncoupling Hsp70 expression from Hsf1 regulation. As predicted by the model, Hsf1 was unable to efficiently deactivate in the absence of Hsp70 transcriptional induction. Next we mapped a discrete Hsp70 binding site on Hsf1 to a motif in the C-terminal activation domain known as conserved element 2 (CE2). Removal of CE2 resulted in increased Hsf1 activity under non-heat shock conditions and delayed deactivation kinetics. In addition, we uncovered a role for the N-terminal domain of Hsf1 in negatively regulating DNA binding. These results reveal the quantitative control mechanisms underlying the feedback loop charged with maintaining cytosolic proteostasis.

## INTRODUCTION

The heat shock response is a transcriptional program conserved in eukaryotes from yeast to humans in which genes encoding molecular chaperones and other components of the protein homeostasis (proteostasis) machinery are activated to counteract proteotoxic stress (Anckar and Sistonen, 2011; Richter et al., 2010). The conserved master transcriptional regulator of the heat shock response, Heat shock factor 1 (Hsf1), binds as a trimer to its cognate DNA motif - the heat shock element (HSE) - in the promoters and enhancers of its target genes (Gross et al., 1990; Hentze et al., 2016; Sorger and Nelson, 1989; Xiao et al., 1991).

In yeast, Hsf1 is essential under all conditions because it is required to drive the high level of basal chaperone expression needed to sustain growth (McDaniel et al., 1989; Solis et al., 2016). Mammalian Hsf1 is dispensable under non-heat shock conditions because it exclusively controls stress-inducible expression of its target regulon, while high-level basal chaperone expression is Hsf1-independent (Mahat et al., 2016). Notably, hsf1^-/-^ mice are not only viable but are in fact resistant to many laboratory cancer models, and Hsf1 has been shown to play procancer roles both in the tumor cells and the supporting stroma (Dai et al., 2012; Dai et al., 2007; Santagata et al., 2011; Scherz-Shouval et al., 2014). In addition to supplying high levels of chaperones to cancer cells, Hsf1 takes on specialized transcriptional roles to support malignant growth, and its activity is associated with poor prognosis in a range of human cancers (Mendillo et al., 2012; Santagata et al., 2011; Scherz-Shouval et al., 2014). Conversely, lack of Hsf1 activity has been proposed to contribute to the development of neurodegenerative diseases associated with protein aggregates (Gomez-Pastor et al., 2017; Neef et al., 2011). Despite the potential therapeutic benefits of modulating Hsf1 activity, a quantitative description of the regulatory mechanisms that control its activity in any cell type remains lacking.

Phosphorylation, SUMOylation, acetylation, chaperone binding (Hsp40, Hsp70, Hsp90 and/or TRiC/CCT), intrinsic thermosensing and an RNA aptamer have all been suggested to regulate Hsf1 in various model systems (Anckar and Sistonen, 2011; Baler et al., 1993; Cotto et al., 1996; Hentze et al., 2016; Hietakangas et al., 2003; Holmberg et al., 2001; Kline and Morimoto, 1997; Neef et al., 2014; Shamovsky et al., 2006; Shi et al., 1998; Westerheide et al., 2009; Xia et al., 1998; Zheng et al., 2016; Zhong et al., 1998; Zou et al., 1998). These diverse mechanisms can operate on Hsf1 by impinging on a number of steps required for activation including nuclear localization, trimerization, DNA binding and recruitment of the transcriptional machinery. Our recent work in the budding yeast *Saccharomyces cerevisiae* demonstrated that binding and dissociation of the chaperone Hsp70 is the primary ON/OFF switch for Hsf1, while phosphorylation is dispensable for activation but serves to amplify the transcriptional output (Zheng et al., 2016).

Based on these results, we generated a mathematical model of the yeast heat shock response. Given that we observed heat shock-dependent dissociation of Hsp70 from Hsf1, and that the genes encoding Hsp70 are major targets of Hsf1, we centered the model on a simple feedback loop in which Hsf1 activates expression of Hsp70, which in turn represses Hsf1 activity. While the model was able to recapitulate experimental data of Hsf1 activity during heat shock and correctly predicted the outcome of defined perturbations, its two central tenets remain untested: 1) Hsp70 directly binds to Hsf1 at a specific regulatory site; 2) Transcriptional induction of Hsp70 provides negative feedback required to deactivate Hsf1. Here, we provide direct evidence supporting these core model assumptions by severing the transcriptional feedback loop, rendering Hsf1 unable to deactivate, and mapping a direct Hsp70 binding site on Hsf1 through which Hsp70 represses its potent C-terminal transactivation domain. These results suggest that the heat shock response circuitry in this model system can be abstracted to a simple two-component feedback loop.

## RESULTS

### Hsp70-mediated negative feedback is required to deactivate Hsf1

Our model of the heat shock response is centered on a feedback loop in which Hsf1 regulates expression of its negative modulator, Hsp70 (Figure 1A). When the temperature is raised, the concentration of unfolded proteins exceeds the capacity of Hsp70. Hsp70 is titrated away from Hsf1, freeing Hsf1 to induce more Hsp70. Once sufficient Hsp70 has been produced to restore proteostasis, Hsp70 binds and deactivates Hsf1. In addition to producing more Hsp70, Hsf1 also induces expression of an inert YFP reporter that can be used as a proxy for Hsf1 activity. In the yeast strains used here, this YFP reporter is integrated into the genome under the control of a promoter containing four repeats of the heat shock cis-element (4xHSE) recognized by Hsf1 (Zheng et al., 2016).

**Figure 1.**
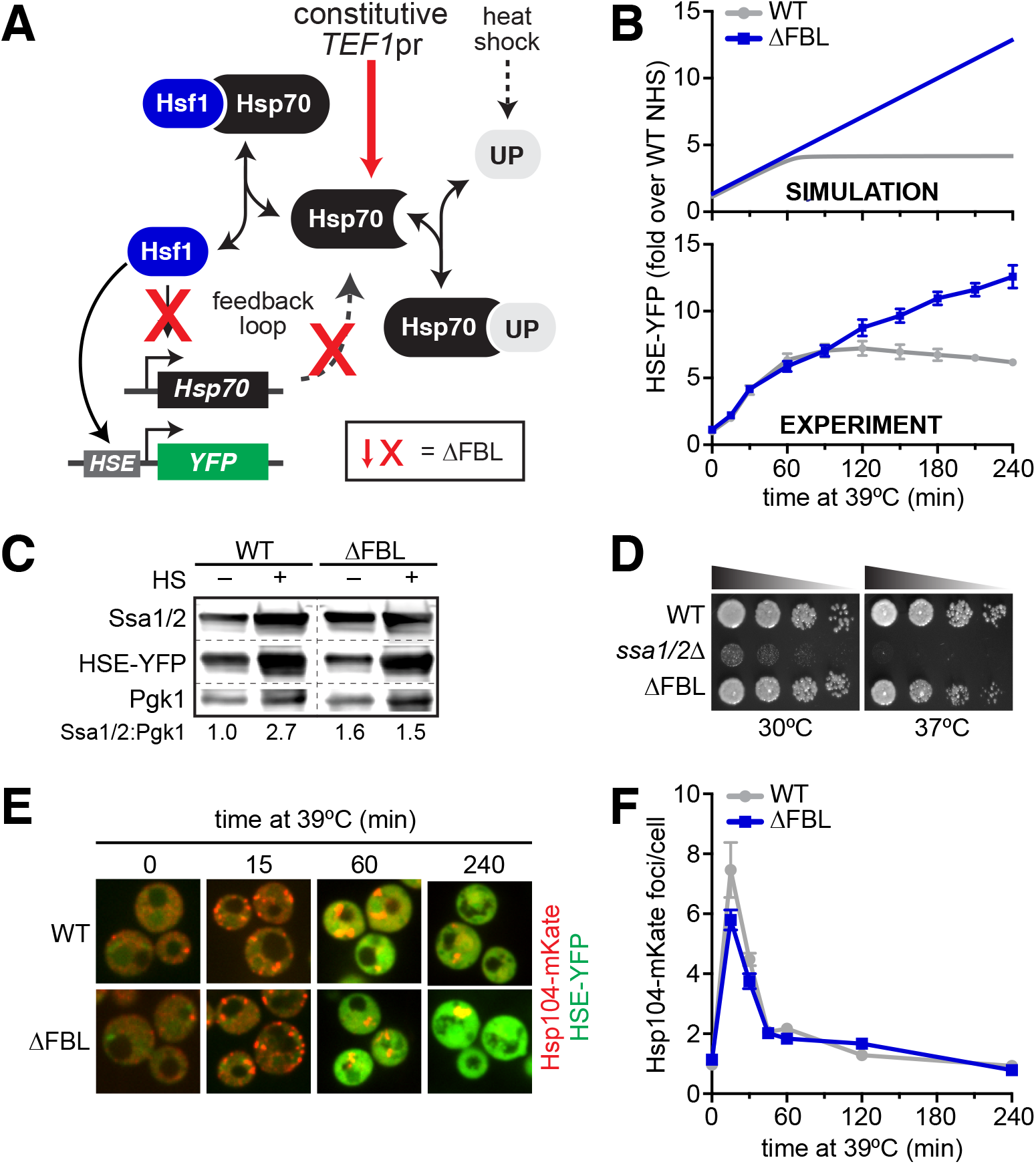
Transcriptional induction of Hsp70 during heat shock is required for Hsf1 deactivation but not proteostasis. **A)** Schematic of the Hsf1 regulatory circuit described by the mathematical model. To generate the feedback-severed yeast strain (ΔFBL), all four Hsp70 paralogs (*SSA1/2/3/4*) were deleted from the genome and 2 copies of *SSA2* under the control of the Hsf1-independent TEF1 promoter were integrated to achieve comparable Hsp70 expression under basal conditions. **B)** Simulated and experimental heat shock time courses comparing the HSE-YFP reporter in wild type and ΔFBL cells. The experimental points represent the average of the median HSE-YFP level in three biological replicates, and the error bars are the standard deviation of the replicates. **C)** Western blot of the expression of Hsp70 (Ssa1/2), the HSE-YFP reporter and glycolytic enzyme Pgk1 in wild type and ΔFBL cells under non-heat shock and heat shock conditions. The dashed lines indicate where lanes were cropped for organization. **D)** Dilution series spot assay of wild type, *ssa1/ΔA* and ΔFBL cells grown at 30°C and 30°C for 36 hours. **E)** Wild type and ΔFBL cells expressing the Hsp104-mKate aggregation reporter along with the HSE-YFP imaged over a heat shock time course showing transient accumulation of Hsp104 foci and sustained induction of HSE-YFP levels in the ΔFBL cells. **F)** Quantification of the number of Hsp104-mKate foci in wild type and ΔFBL cells over a heat shock time course. N > 100 cells for each time point. Error bars represent the standard error of the mean.

To test the model, we severed the feedback loop, both computationally and experimentally, and monitored Hsf1 activity over time following a shift from 25ºC to 39ºC by simulating and measuring the HSE-YFP reporter. We cut the feedback loop in the mathematical model by removing the equation relating the production of Hsp70 to the concentration of free Hsf1 without changing any parameters or initial conditions. In the absence of Hsf1-dependent transcription of Hsp70, the model predicted that the HSE-YFP reporter should be activated with the same kinetics as that of the wild type, but should continue to accumulate long after the response is attenuated in the wild type (Figure 1B).

To experimentally test this in yeast cells, we decoupled expression of all four cytosolic Hsp70 paralogs (*SSA1/2/3/4*) from Hsf1 regulation while maintaining the expression of total Hsp70 at its endogenous levels under non-heat shock conditions. This was achieved by integrating two copies of *SSA2* under the control of the Hsf1-independent *TEF1* promoter into the genome and deleting *ssa1/2/3/4.* We named this strain ΔFBL to denote that we had removed the feedback loop (Figure 1A). As expected, wild type cells were able to increase Hsp70 levels and induce the HSE-YFP reporter protein during heat shock, while ΔFBL cells were only able to induce the HSE-YFP protein - but not Hsp70 - during heat shock (Figure 1C). We performed a heat shock time course in WT and ΔFBL cells and compared the expression of the HSE-YFP reporter by flow cytometry. As predicted by the simulation, the ΔFBL strain activated the reporter with identical kinetics to the wild type during the early phase of the response, but failed to attenuate induction during prolonged exposure to elevated temperature (Figure 1B). While the simulation correctly predicted the experimental results qualitatively, the model underestimated the amount of time required to observe the separation between the wild type and ΔFBL strains, suggesting the strength of the feedback had been exaggerated in the first iteration of the model. By reducing the strength of the feedback loop, we were able to quantitatively match the behavior of both the wild type and ΔFBL cells (Figure 1B, see methods for updated parameter values).

The inability of Hsf1 to deactivate in the ΔFBL strain could result either from a specific disruption of the "OFF switch” or from a general failure of the cells to restore proteostasis. In other words, does cutting the feedback loop simply result in sustained stress, or is the prolonged Hsf1 activity the result of specifically breaking its deactivation mechanism? To distinguish these possibilities, we first compared growth of wild type, ΔFBL and *ssa1/2Δ* cells at 30°C and 37°C. The *ssa1/2Δ* cells - which retain viability due to Hsf1-mediated induction of *SSA3/4* - displayed severely impaired growth at 30°C and were inviable at 37°C (Figure 1D). By contrast, the wild type and ΔFBL strains grew equally at 30°C, and the ΔFBL strain showed only a slight reduction in growth at 37°C (Figure 1C). The reduced growth of the ΔFBL mutant at 37°C could be a consequence of either an inadequate or overzealous heat shock response, and does not necessarily indicate a general failure to restore proteostasis. To directly monitor the loss and restoration of proteostasis, we imaged wild type and ΔFBL cells expressing Hsp104-mKate over a heat shock time course. Hsp104 is a disaggregase that forms puncta marking protein aggregates when tagged with a fluorescent protein. Upon acute heat shock, the number of Hsp104-mKate foci spiked in both wild type and ΔFBL cells, but dissolved with the same kinetics in both strains (Figure 1E, F). These data indicate that the ΔFBL cells can restore proteostasis just as efficiently as wild type cells, and suggest that the prolonged Hsf1 activation in the ΔFBL cells is due to a deactivation defect. Thus, the transcriptional negative feedback loop is required to deactivate Hsf1 once proteostasis has been restored.

### Scanning mutagenesis reveals three independent repressive segments in Hsf1

In addition to positioning the transcriptional feedback loop as the core regulatory circuit that controls Hsf1 activity, the model also posits that Hsp70 binding is the mechanism that represses Hsf1. If this assumption is true, then disrupting the binding interaction should increase Hsf1 activity under non-heat shock conditions (Figure 2 - figure supplement 1). To test this, we generated a series of 48 Hsf1 mutants in which we systematically removed 12 amino acid segments along the nonessential N- and C-terminal regions of Hsf1 (Figure 2A). We integrated these mutants into the genome as the only copy of *HSF1* in a strain background bearing the HSE-YFP reporter and assayed for activity by measuring YFP levels under non-heat shock and heat shock conditions by flow cytometry (Zheng et al., 2016). To benchmark the assays, we used wild type Hsf1 and mutants lacking the entire N- and C-terminal regions. As previously shown, removal of the N-terminal region led to significantly increased Hsf1 activity under both non-heat shock and heat shock conditions in this assay (Sorger, 1990; Zheng et al., 2016), while removal of the C-terminal region significantly reduced Hsf1 activity under both conditions (Figure 2A). In the N-terminal region, we found two distinct 12 amino acid segments that when deleted resulted in increased Hsf1 activity (amino acids 85-96 and 121-132) (Figure 2A). In the C-terminal region, removal of two consecutive 12 amino acid segments as well as truncation of the final 6 amino acids resulted in increased Hsf1 activity (amino acids 528-539, 540-551 and 828-833) (Figure 2A).

**Figure 2.**
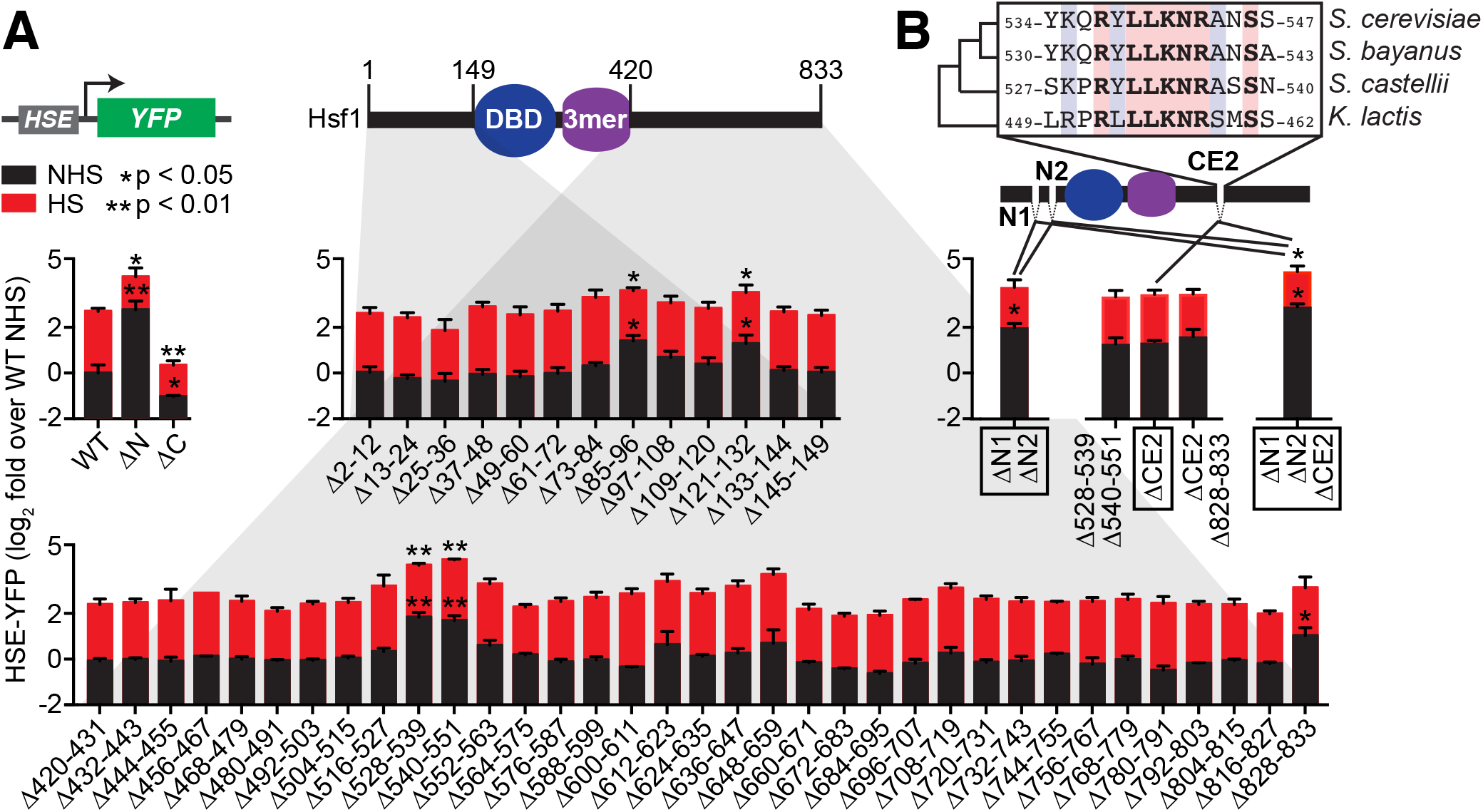
Identification of negative regulatory determinants in the N- and C-termini of Hsf1. **A)** Screen for functional determinants. The indicated Hsf1 mutants were integrated into the genome as the only copy of Hsf1 expressed from the endogenous *HSF1* promoter in a strain expressing the HSE-YFP reporter. Hsf1ΔN is a deletion of the first 145 amino acids following the methionine; Hsf1ΔC is a truncation of the last 409 amino acids of Hsf1, retaining the first 424 amino acids. Each mutant in the scanning deletion analysis is missing a stretch of 12 amino acids in either the N-terminal 149 residues or final 414 C-terminal residues. Each strain was assayed in triplicate for its HSE-YFP level under non-heat shock (NHS) and heat shock (HS) conditions by flow cytometry. The error bars are the standard deviation of the replicates. Statistical significance was determined by a two-tailed T-test (* p < 0.05; ** p < 0.01). **B)** Analysis of double and triple mutants of the functional segments. ΔN1 and ΔN2 represent Δ85-96 and Δ121-132, respectively, and each independently contribute to Hsf1 activity. CE2 is a region spanning the consecutive C-terminal determinants defined in **(A)** that is conserved among a subset of fungal species. Statistical significance was determined by two-tailed T-tests comparing each double mutants to both of the single mutant parents (* p < 0.05 for both T-tests).

To determine if these segments acted independently, we generated double mutants. Combining the N-terminal deletions (Δ85-96/Δ121-132) resulted in a mutant with significantly greater basal activity than either of the single mutants, suggesting that these segments operate independently to repress Hsf1 activity (p < 0.05, Figure 2B). We will refer to these N-terminal segments as N1 and N2. By contrast, combining the consecutive C-terminal segments (Δ528-539/Δ540-551) resulted in a double mutant with the same activity as the single deletions, suggesting that a unique functional determinant encompasses these segments (Figure 2B). Consistent with this notion, a region spanning these two segments comprises a previously identified element conserved in Hsf1 in other fungal species known as the "conserved element 2” (CE2) (Figure 2B) (Jakobsen and Pelham, 1991). Indeed, specific removal of CE2 was sufficient to match the increased level of Hsf1 activity observed in the Δ528-539/Δ540-551 mutant (Figure 2B). Additional removal of the final 6 amino acids provided no further increase in Hsf1 activity, consistent with previous studies suggesting a non-additive interaction between these elements (Figure 2B) (Hashikawa and Sakurai, 2004; Yamamoto et al., 2007). However, combining the N1/N2 and CE2 deletions resulted in an Hsf1 mutant with significantly greater activity than either the ΔN1/ΔN2 mutant or the ΔCE2 mutant (Figure 2B). Together, the scanning mutagenesis revealed three independent repressive segments on Hsf1 (N1, N2, and CE2).

### N1/N2 regulate DNA binding while CE2 regulates transactivation

The segments we identified with increased HSE-YFP levels could function either by enhancing the association of Hsf1 with HSEs (i.e., increasing DNA binding) or by boosting the transactivation capacity of Hsf1 (i.e., increasing recruitment of the transcriptional machinery). To directly test the ability to bind to HSEs in cells, we performed chromatin immunoprecipitation (ChIP) of wild type Hsf1, Hsf1^ΔN^, Hsf1^ΔC^, Hsf1^ΔN1/ΔN2^, Hsf1^ΔCE2^ and Hsf1^ΔN1/ΔN2/ΔCE2^ under non-heat shock and acute (5 minute) heat shock conditions. Following ChIP enrichment, we assayed for association with the synthetic *4xHSE* promoter that drives the YFP reporter as well as five endogenous target gene promoters (*HSC82, HSP82, SSA4, HSP26* and *TMA10*) by qPCR. Under non-heat shock conditions, wild type Hsf1 binding ranged over nearly two orders of magnitude across these targets, from 0.14% of input at the *TMA10* promoter to 12.0% of input at the *4xHSE* promoter (Figure 3—figure supplement 1). Upon acute heat shock, the inducibility of Hsf1 binding also varied widely across these targets, with induction of greater than 100-fold for *TMA10* and less than 1.5-fold for *HSC82* (Figure 3—figure supplement 1). These data are inconsistent with the notion that Hsf1 is constitutively bound to its target genes (Jakobsen and Pelham, 1988; Sorger et al., 1987).

Interestingly, both the Hsf1^ΔN^ and Hsf1^ΔC^ mutants showed significantly increased association with the *4xHSE* and *SSA4* promoters under non-heat shock conditions (Figure 3A, Figure 3— figure supplement 1). However, while increased binding to the *4xHSE* promoter was accompanied by increased transcriptional output of the YFP reporter in Hsf1^ΔN^ cells, no such increase in HSE-YFP levels was observed in Hsf1^ΔC^ cells (Figure 3B). In fact, Hsf1^ΔC^ cells showed significantly reduced HSE-YFP levels under non-heat shock conditions compared to wild type (Figure 2A). These data suggest a simple relationship between DNA binding and transcription for the Hsf1^ΔN^ mutant: the N-terminal region of Hsf1 inhibits DNA binding and thereby reduces transcriptional activity. By contrast, there is no correlation between DNA binding and transcription for the Hsf1^ΔC^ mutant.

**Figure 3.**
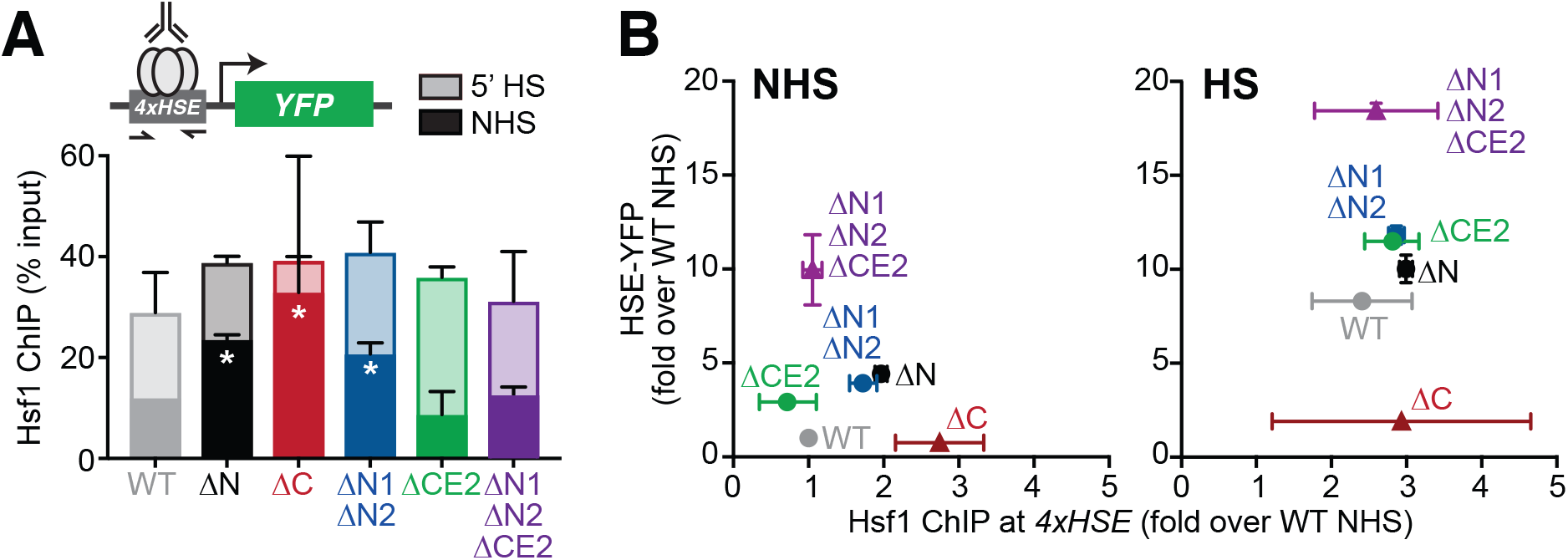
The Hsf1 N-terminus regulates DNA binding while CE2 controls transactivation. **A)** Chromatin immunoprecipitation of Hsf1 followed by quantitative PCR of the *4xHSE* promoter in the indicated Hsf1 wild type and mutant strains under non-heat shock and heat shock conditions (solid and outlined bars, respectively). Error bars show the standard deviation of biological replicates. Statistical significance was determined by a two-tailed T-test (* p < 0.05; ** p < 0.01). **B)** Relationship between Hsf1 binding at the 4xHSE promoter as determined by ChIP-qPCR and transcriptional activity as measured by levels of the HSE-YFP reporter under non-heat shock (NHS) and heat shock (HS) conditions for the panel of mutants assayed in **(A)**.

Consistent with a role for the N-terminal segment in regulating DNA binding, the Hsf1^ΔN1/ΔN2^ mutant mirrored Hsf1^ΔN^ in both its increased binding to the *4xHSE* promoter and increased transcription of the YFP reporter under non-heat shock conditions relative to wild type (Figure 3A, B). However, unlike the complete ablation of the N-terminal region, Hsf1^ΔN1/ΔN2^ showed no increase in association with the *SSA4* promoter compared to wild type (Figure 3—figure supplement 1), suggesting that its enhanced association with endogenous targets may be limited. Neither Hsf1^ΔCE2^ nor Hsf1^ΔN1/ΔN2/ΔCE2^ showed any significant differences compared to wild type at any of the six target promoters under either non-heat shock or heat shock conditions, indicating that CE2 has no effect on Hsf1 DNA binding (Figure 3—figure supplement 1). Remarkably, under heat shock conditions, none of the five mutants showed significant differences in binding to the *4xHSE* promoter compared to wild type (Figure 3A). Thus, during heat shock, the differences in YFP reporter levels reflect the different transactivation abilities of the series of mutants, spanning more than 16-fold between Hsf1^ΔC^ and Hsf1^ΔN1/ΔN2/ΔCE2^ (Figure 3B). Taken together, the ChIP results suggest that multiple determinants, including the N1/N2 segments and the C-terminal domain, contribute to regulating DNA binding.

### CE2 is a direct binding site for Hsp70

Since CE2 affects Hsf1 transactivation but not DNA binding, we hypothesized that it could be a binding site for Hsp70. To test this, we performed serial immunoprecipitation from whole cell lysates followed by mass spectrometry (IP/MS) of 3xFLAG/V5-tagged Hsf1 mutants to identify specific interactions with chaperone proteins. We measured Hsp70 binding to wild type Hsf1, Hsf1^ΔN^, Hsf1^ΔC^, Hsf1^ΔN1/ΔN2^, Hsf1^ΔCE2^ and Hsf1^ΔN1/ΔN2/ΔCE2^ under non-heat shock conditions, performing three biological replicates for each. Removal of the entire N-terminal region or the N1/N2 segments had no effect on Hsp70 binding relative to wild type, consistent with a role confined to regulating DNA binding (Figure 4A). By contrast, removal of the full C-terminal region significantly reduced the association of Hsf1 with Hsp70 (Figure 4A). Moreover, specific removal of CE2 - either alone or in combination with the N1/N2 deletions - also resulted in significantly diminished association with Hsp70, nearly matching removal of the entire C-terminal region (Figure 4A). Analysis of an additional biological replicate by Western blotting corroborated the IP/MS results (Figure 4A). The residual Hsp70 that co-precipitated with Hsf1^ΔCE2^ was refractory to dissociation upon heat shock, suggesting that this secondary interaction is unlikely to be regulatory (Figure 4B).

**Figure 4.**
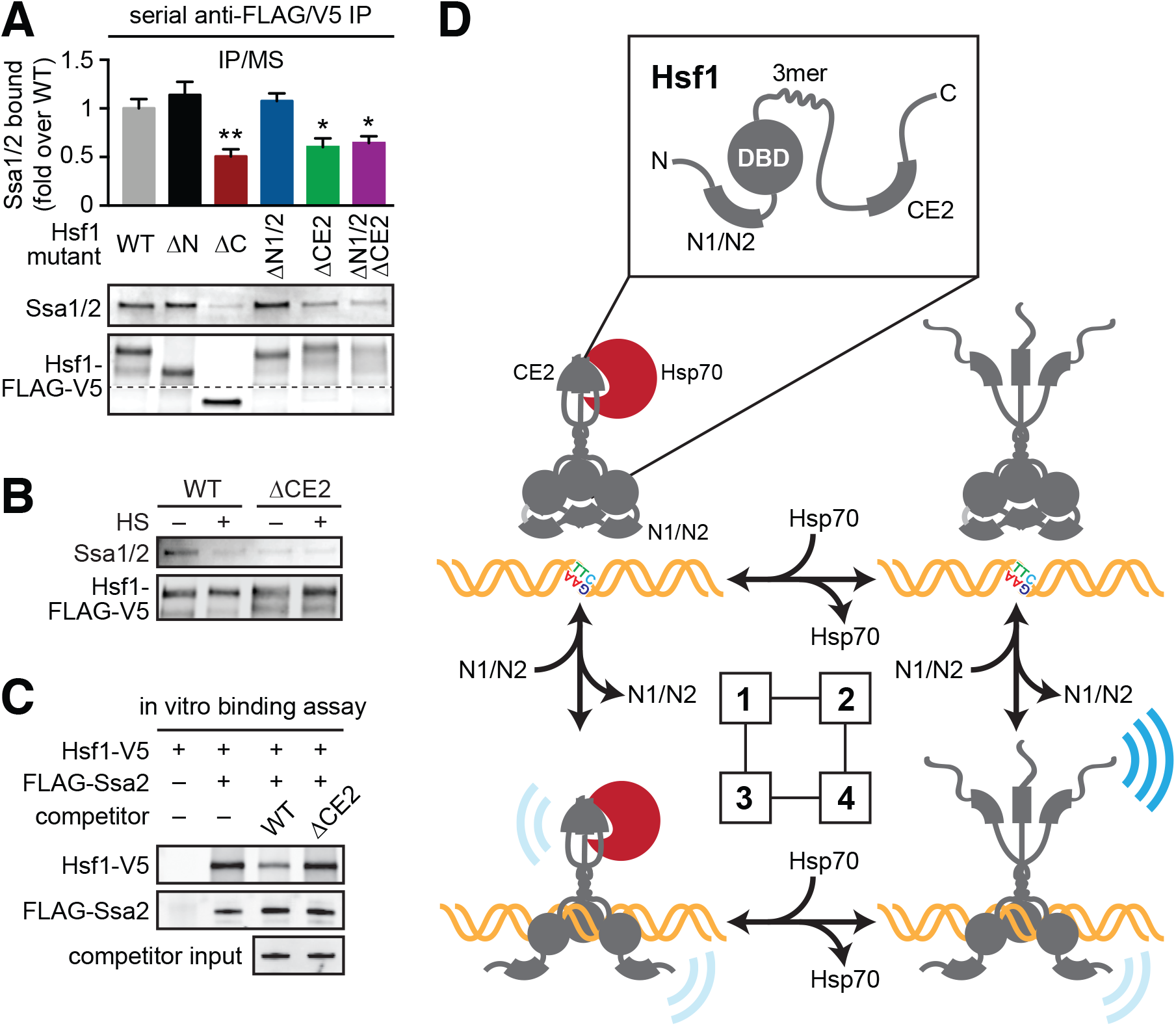
CE2 is a direct Hsp70 binding site. **A)** Co-immunoprecipitation of Hsf1 and Hsp70. The indicated Hsf1 mutants, C-terminally tagged with 3xFLAG-V5, were serially precipitated and subjected to mass spectrometry as described. The ratio of Hsp70 (Ssa1/2) to Hsf1 was determined in three three biological replicates (bar graph, error bars are the standard deviation). Statistical significance was determined by a twotailed T-test (* p < 0.05; ** p < 0.01). An additional replicate was analyzed by Western blot using antibodies against Ssa1/2 and the FLAG tag to recognize Hsf1. The FLAG blot was cropped in the middle to show the much smaller Hsf1^ΔC^. **B)** Cells expressing C-terminally 3xFLAG-V5-tagged wild type Hsf1 and Hsf1^ΔCE2^ were either left untreated or heat shocked for 5 minutes at 39°C before serial Hsf1 imunnoprecipitation and analyzed by Western blot using antibodies against Ssa1/2 and the FLAG tag to recognize Hsf1. **C)** In vitro Hsf1:Hsp70 binding assay. Recombinant Hsf1-V5 and 3xFLAG-Ssa2 were purified, incubated together and assayed for binding by anti-FLAG immunoprecipitation followed by epitope-tag-specific Western blot. Addition of 5-fold molar excess of wild type Hsf1-6xHIS but not Hsf1^ΔCE2^-6xHIS diminished the amount of Hsf1-V5 bound to 3xFLAG-Ssa2. **D)** Thermodynamic representation of the 4 state model of Hsf1 activity.

Simulations of heat shock time courses as a function of decreased affinity between Hsf1 and Hsp70 predicted progressively increased levels of the HSE-YFP under non-heat shock conditions and prolonged activation following heat shock relative to wild type (Figure 4—figure supplement 1A). In agreement, Hsf1^ΔCE2^ has elevated HSE-YFP levels under non-heat shock conditions and displayed delayed deactivation kinetics compared to wild type in a heat shock time course (Figure 4—figure supplement 1B).

Finally, to test a direct role for CE2 in binding to Hsp70, we utilized an in vitro binding assay we previously established to monitor interaction between recombinant purified Hsf1 and Hsp70 (Zheng et al., 2016). Whereas wild type Hsf1-6xHIS was able to outcompete wild type Hsf1-V5 for binding to the Hsp70 Ssa2 at a 5-fold molar excess, Hsf1^ΔCE2^-6xHIS was not (Figure 4C). These results demonstrate that CE2 is a direct binding site for Hsp70 through which Hsp70 represses Hsf1.

## DISCUSSION

In this study we tested the assumptions of our mathematical model of the heat shock response by severing the Hsp70 transcriptional feedback loop and mapping an Hsp70 binding site on Hsf1. While we uncovered more biological complexity in Hsf1 regulation than we represent in the model, we validated the model’s central tenets - that Hsp70 binding and dissociation turn Hsf1 off and on, and that transcriptional induction of Hsp70 represents a critical negative feedback loop required for the homeostatic regulation of Hsf1. Moreover, we found the model to be remarkably powerful in its ability to predict the dynamics of Hsf1 activity when challenged with targeted perturbations to the system architecture despite its oversimplified structure. These results argue that conceptualizing the heat shock response as a two-component feedback loop - in which Hsf1 positively regulates Hsp70 expression and Hsp70 negatively regulates Hsf1 activity - is an appropriate abstraction that captures the essence of the regulatory network. Whether this simplifying abstraction can be applied to HSF1 regulation in metazoans remains to be determined.

At a more mechanistic level, our screen for functional determinants in the N- and C-terminal regions of Hsf1 revealed three distinct segments in Hsf1 that independently exert negative regulation. The two N-terminal segments contribute to hitherto unknown repression of Hsf1 DNA binding, while the single C-terminal segment, CE2, is a binding site for Hsp70 through which Hsp70 represses Hsf1 transactivation. Although, as its name suggests, CE2 is conserved, it is restricted to a subset of yeast species and is absent in mammalian HSF1 sequences. Its amino acid composition, consisting of hydrophobic and basic residues, is reminiscent of peptide sequences known to bind to Hsp70 in vitro (Van Durme et al., 2009), lending additional credence to our results. Thus, while CE2 is not conserved in mammalian genomes in primary sequence, it would seem facile to evolve a distinct but functionally analogous hydrophobic and basic segment to allow for Hsp70 binding. Notably, even though we found no evidence that the N1 segment is an additional Hsp70 binding site on endogenous Hsf1, its sequence is also predicted to be an Hsp70 binding site and is capable of binding to Hsp70 when overexpressed *in trans* (S. Peffer and K. Morano, personal communication).

In addition to mechanistic insight into Hsp70 binding, our results for the first time reveal the existence of intramolecular determinants that negatively regulate Hsf1 DNA binding. While it has been known for many years that removal of the N-terminal region of Hsf1 leads to increased activity (Sorger, 1990) - suggesting that this region is repressive in nature - the N-terminus also has a transactivation function and is important for efficient recruitment of Mediator during heat shock (Kim and Gross, 2013). Here we show that removal of the full N-terminal region results in increased association with target gene promoters under non-heat shock conditions (Figure 3A), indicating a role for this yeast-specific region in impeding DNA binding and suggesting a mechanistic basis for the increased transcriptional activity of Hsf1^ΔN^ relative to wild type Hsf1. In particular, the N1/N2 segments contribute to blocking DNA binding, as Hsf1^ΔN/ΔN2^ displayed increased association with the synthetic *4xHSE* promoter (Figure 3A). If N1 were a bona fide second Hsp70 binding site (Peffer and Morano, personal communication), then Hsp70 would be likely to regulate both Hsf1 DNA binding and transactivation. Alternatively, if the N1/N2 segments impede DNA binding independent of Hsp70, then an additional heat shock-dependent mechanism would be required to relieve this block. Perhaps, by analogy to the intrinsic ability of human HSF1 to trimerize and bind DNA at elevated temperature (Hentze et al., 2016), the N1/N2 segments could contribute to direct thermosensing by mediating a temperature-dependent conformational change that increases DNA binding ability. The role of the C-terminus in regulating Hsf1 DNA binding is less clear, given that we observed increased association with the 4xHSE promoter yet diminished HSE-YFP levels. There could be an element in the C-terminus that inhibits Hsf1 DNA binding. Alternatively, the increased DNA association observed for Hsf1^ΔC^ could be a consequence of its severely impaired transactivation ability: If each binding event is less likely to lead to productive transcription, then the cell must force Hsf1^ΔC^ to compensate to achieve sufficient transcription of the essential Hsf1 regulon; thus, Hsf1^ΔC^ must engage in more binding events to sustain growth. Moreover, since Hsf1^ΔC^ has to use its N-terminal region as a transactivator, the N-terminus may be unavailable to impede DNA binding.

Putting all of these observations together, we propose that Hsf1 can exist in one of four states in the yeast nucleus (Figure 4D):

1) *C-terminal activation domain (CTA) closed/DBD unbound* Hsp70 is bound to CE2 keeping the CTA closed; the N-terminal region is engaged in blocking the DBD from accessing available HSEs via the N1/N2 segments.
2) *CTA open/DBD unbound* Hsp70 has dissociated from CE2; the CTA is open and can potentially recruit the transcriptional machinery; the N-terminal region continues to hinder DNA binding.
3) *CTA closed/DBD bound* Hsp70 remains bound to CE2 keeping the CTA closed; the N-terminal region has reoriented to allow HSE binding; Hsf1 weakly recruits the transcriptional machinery.
1) *CTA open/DBD bound* Hsp70 has dissociated from CE2 and the CTA is open; the N-terminal region has reoriented to allow HSE binding; Hsf1 avidly recruits the transcriptional machinery.

The dual mechanisms of Hsf1 regulation described here - control of DNA binding and accessibility of the transactivation domain - in combination with the fine-tuning capacity we previously demonstrated for phosphorylation (Zheng et al., 2016), exert exquisite quantitative control over the Hsf1 regulon. We propose that these three regulatory mechanisms enable cells to precisely tailor an optimal response to a variety of environmental and internal stresses.

## ACKNOWLEDGEMENTS

We are grateful A. Kane for providing us with the phleomycin resistance cassette and deleting SSA3, to A. Jaeger for beneficial discussions, and to H. Lodish, G. Fink and their lab members for insightful comments. Experimentally, we are indebted to E. Spooner and the Whitehead Proteomics core for mass spectrometric analysis, to the Whitehead Institute FACS facility for technical assistance and to N. Azubuine and T. Nanchung for a constant supply of plates and media. This work was supported by an NIH Early Independence Award (DP5 OD017941-01 to D.P.), a National Science Foundation CAREER Award (MCB-1350949 to A.S.K.) and a National Science Foundation grant (MCB-1518345 to D.S.G.).

## AUTHOR CONTRIBUTIONS

Conceptualization, D.P. and A.S.K; Methodology, J.K., X.Z., N.P., K.V., A.J. and D.P.; Investigation, J.K., X.Z., N.P., A.J., K.V., and D.P.; Writing, D.P., D.S.G. and A.S.K.; Funding Acquisition, D.P., A.S.K. and D.S.G.; Supervision, D.P., A.S.K. and D.S.G.

## METHODS

### Yeast strains, plasmids and cell growth

Yeast cells were cultured in SDC media and dilution series spot assays were performed as described. Strains and plasmids are listed in Supplementary Tables 1 and 2.

### Mathematical modleling

Modeling was performed as described (Zheng et al., 2016).

Model parameter:

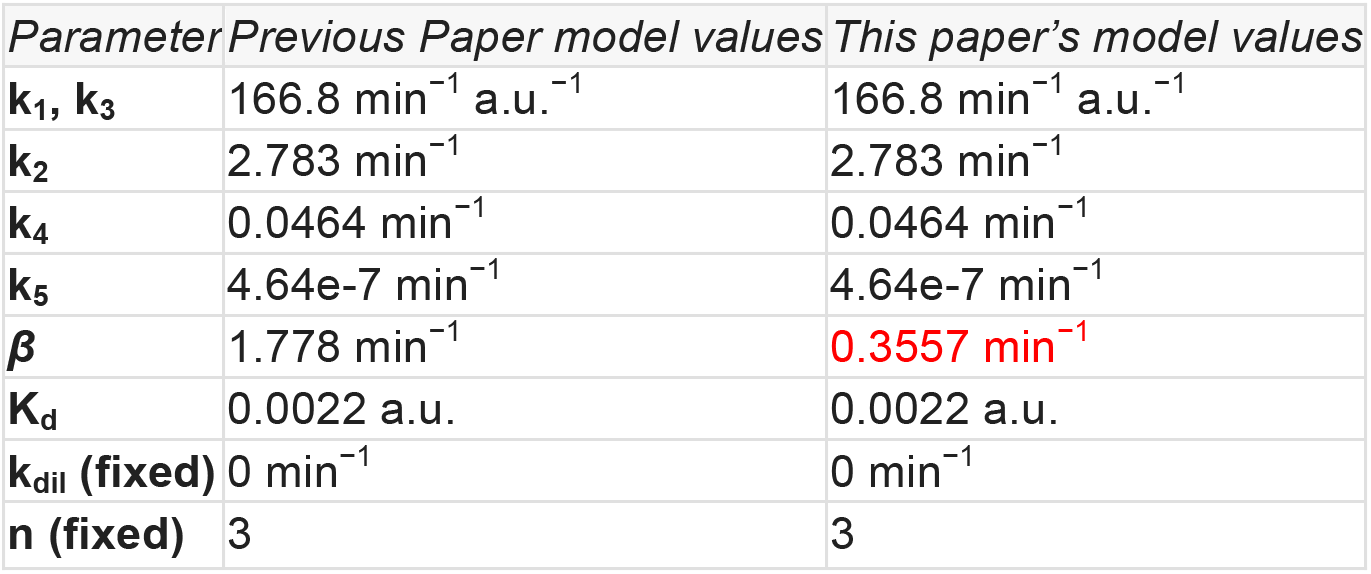

| *Parameter* | *Previous Paper model values* | *This paper's model values* |
| --- | --- | --- |
| $k_1, k_3$ | $166.8 \text{ min}^{-1} \text{ a.u.}^{-1}$ | $166.8 \text{ min}^{-1} \text{ a.u.}^{-1}$ |
| $k_2$ | $2.783 \text{ min}^{-1}$ | $2.783 \text{ min}^{-1}$ |
| $k_4$ | $0.0464 \text{ min}^{-1}$ | $0.0464 \text{ min}^{-1}$ |
| $k_5$ | $4.64\text{e-}7 \text{ min}^{-1}$ | $4.64\text{e-}7 \text{ min}^{-1}$ |
| $\beta$ | $1.778 \text{ min}^{-1}$ | $0.3557 \text{ min}^{-1}$ |
| $K_d$ | 0.0022 a.u. | 0.0022 a.u. |
| $k_{dil}$ (fixed) | $0 \text{ min}^{-1}$ | $0 \text{ min}^{-1}$ |
| $n$ (fixed) | 3 | 3 |

Initial conditions:

| *Species* | *Initial value (a.u.)* | *Description* |
| --- | --- | --- |
| [HSP]<sub>o</sub> | 1 | Free Hsp70 |
| [Hsf1]<sub>o</sub> | 0 | Free Hsf1 |
| [HSP•Hsf1]<sub>o</sub> | 0.002 | HSP70•Hsf1 complex |
| [HSP•UP]<sub>o</sub> | 0 | Hsp70•UP complex |
| [YFP]<sub>o</sub> | 3 | Initial YFP concentration |
| [UP]<sub>o</sub> (@ 39°C) | 10.51 | UP concentration at 39°C |

### Flow cytometry

Heat shock experiments and heat shock time courses were performed and HSE-YFP levels were quantified by flow cytometry as described (Zheng et al., 2016). Data were processed in FlowJo 10. Data were left ungated and YFP fluorescence was normalized by side scatter (SSC) for each cell.

### Spinning disc confocal imaging

Imaging was performed as described (Zheng et al., 2016). Hsp104-mKate foci were quantified manually in ImageJ.

### Chromatin Immunoprecipitation (ChIP)

Hsf1 ChIP was performed and quantified by qPCR as described (Anandhakumar et al., 2016).

### Serial 3xFLAG/V5 immunoprecipation

Hsf1-3xFLAG-V5 was serially immunoprecipitated and analyzed by mass spectrometry and Western blotting as described (Zheng et al., 2016; Zheng and Pincus, 2017).

### Recombinant protein binding and competition assay

In vitro binding assay between Hsf1 and Ssa2 was performed as described (Zheng et al., 2016).

---

**Figure 2—figure supplement 1.**
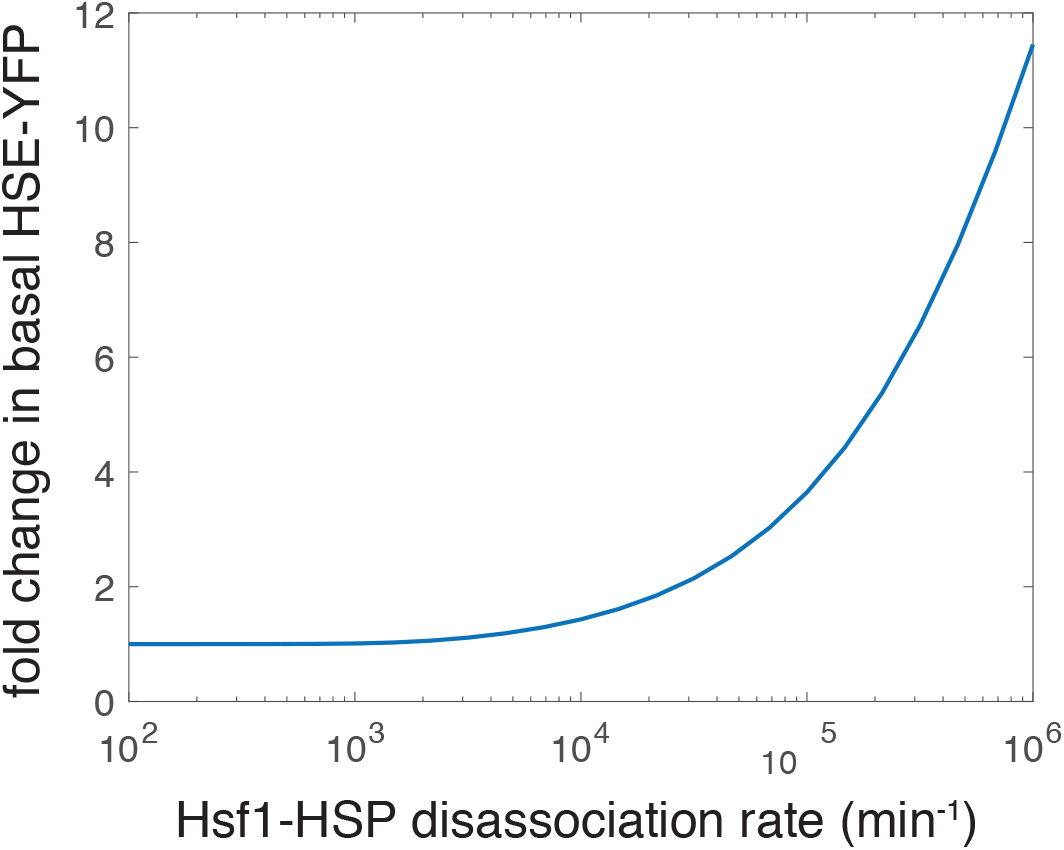
Simulation showing an increase in the basal level of the HSE-YFP reporter as a function of increased dissociation rate (decreased affinity) of the Hsp70-Hsf1 interaction. The "wild type” rate is 2.783 min^-1^ as in the previous iteration of the model (not shown on the graph) (Zheng et al., 2016).

**Figure 3—figure supplement 1.**
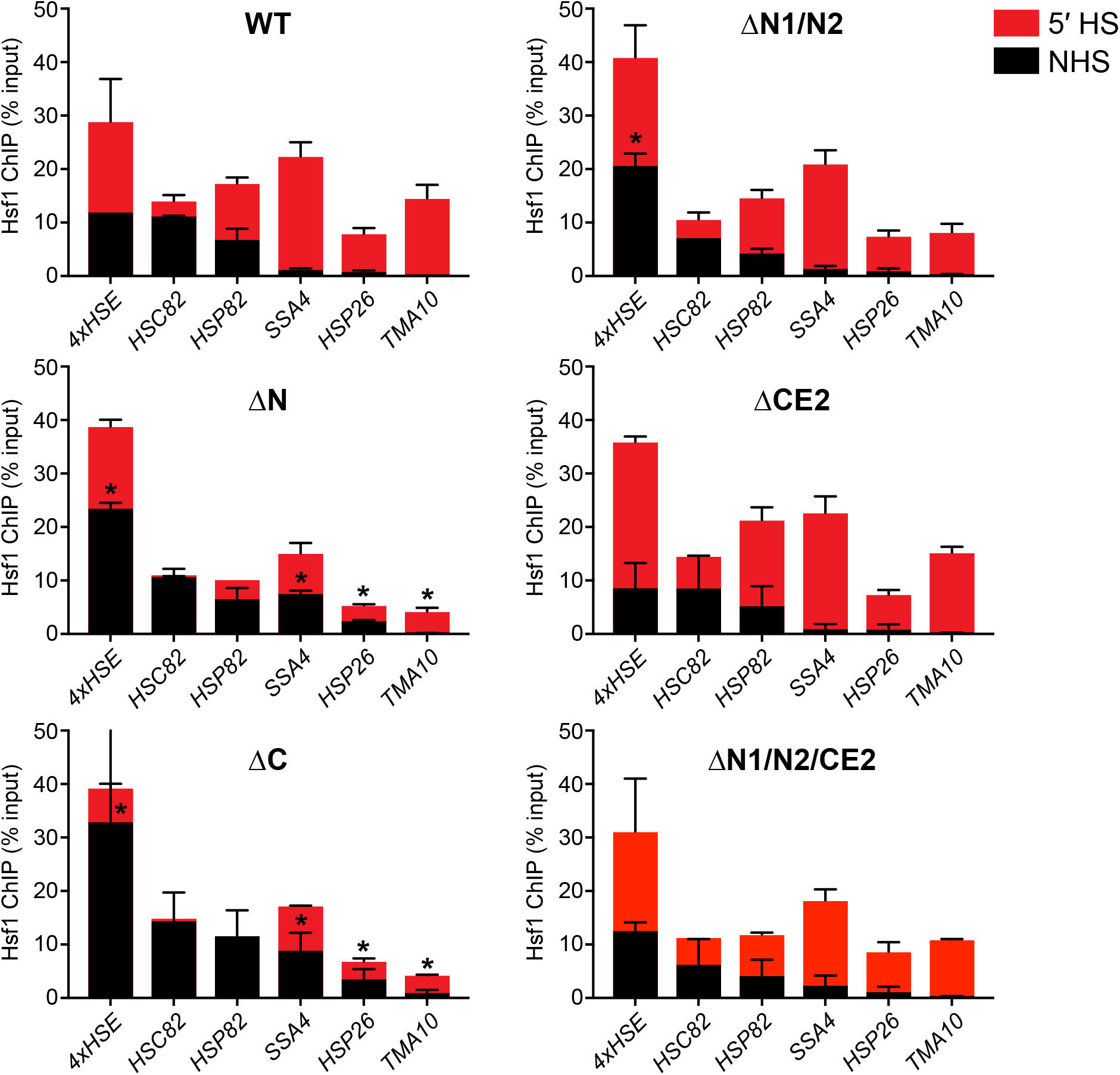
ChIP-qPCR of Hsf1 mutants at endogenous target promoters under non-heat shock and heat shock conditions. Error bars show the standard deviation of biological replicates. Statistical significance was determined by a two-tailed T-test (* p < 0.05; ** p < 0.01).

**Figure 4—figure supplement 1.**
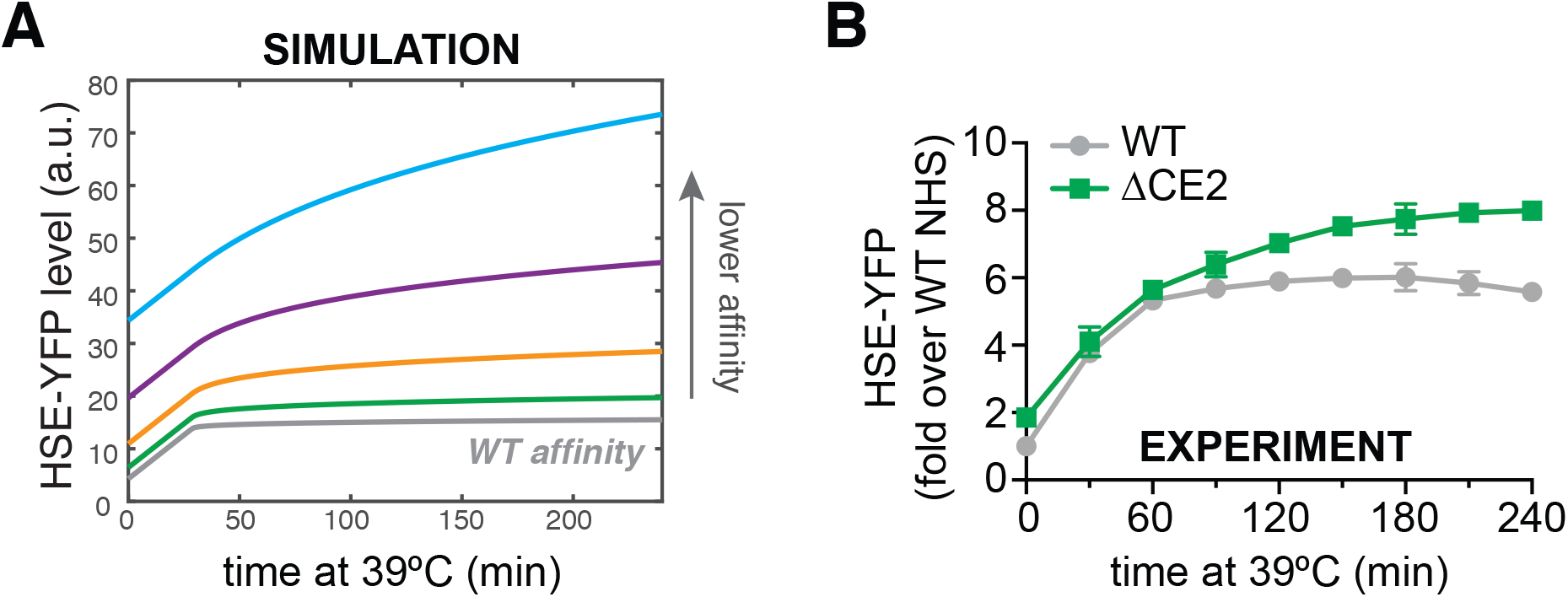
Reduced affinity for Hsp70 results in increased basal Hsf1 activity and delayed deactivation kinetics during heat shock. **A)** Simulations of HSE-YFP levels over a heat shock time course as a function of increased rate of dissocation (reduced affinity) of Hsp70 from Hsf1. **B)** Experimental heat shock time course of HSE-YFP levels in cells expressing wild type Hsf1 or Hsf1^ΔCE2^. Each point represents the average of the median HSE-YFP level in three biological replicates, and the error bars are the standard deviation of the replicates.

